# Ischemic stroke increases levels of the folate receptor and one-carbon enzymes in male and female brain tissue

**DOI:** 10.1101/2025.09.02.673861

**Authors:** Petter Burrows, Himmat Dhillon, Gillian E. McDemott, Amanda Covaleski, Lilah Manfredi, Thomas G Beach, Geidy E Serrano, Nafisa M. Jadavji

## Abstract

Stroke is the second most common cause of death worldwide and predominantly affects individuals over 65 years old. Its prevalence is projected to increase in parallel with the aging global population. Nutrition is a modifiable risk factor for ischemic stroke. Folates, B-vitamins and choline play a central role in one-carbon metabolism (1C), which is a key metabolic network that integrates nutritional signals with biosynthesis, redox homeostasis, epigenetics, regulation of cell proliferation, and stress resistance. Our research group has previously shown that deficiencies in 1C lead to worsened stroke outcomes using preclinical models. However, the impact of ischemic stroke on 1C enzymes in affected brain tissue remains unknown. The objective of this study is to investigate whether ischemic stroke contributes to a change in the levels of 1C enzymes after ischemic stroke in male and female patients. Cortical brain tissue sections from ischemic stroke patients and controls were stained for enzymes involved in 1C. All tissue was co-stained with neuronal nuclei (NeuN) and DAPI (4′,6-diamidino-2-phenylindole). The colocalization of all three markers was evaluated by two individuals who were blinded to the experimental groups. Ischemic stroke increased neuronal levels of the folate receptor and 1C enzymes, methylenetetrahydrofolate reductase (MTHFR), thymidylate synthase (TS) and serine hydroxy methyltransferase (SHMT). In male stroke brain tissue was observed to have increased levels of MTHFR, TS, and SHMT. Female brain tissue had increases in the folate receptor and TS. The results suggest that ischemic stroke leads to increased demand of 1C and that there are some differences between males and females.

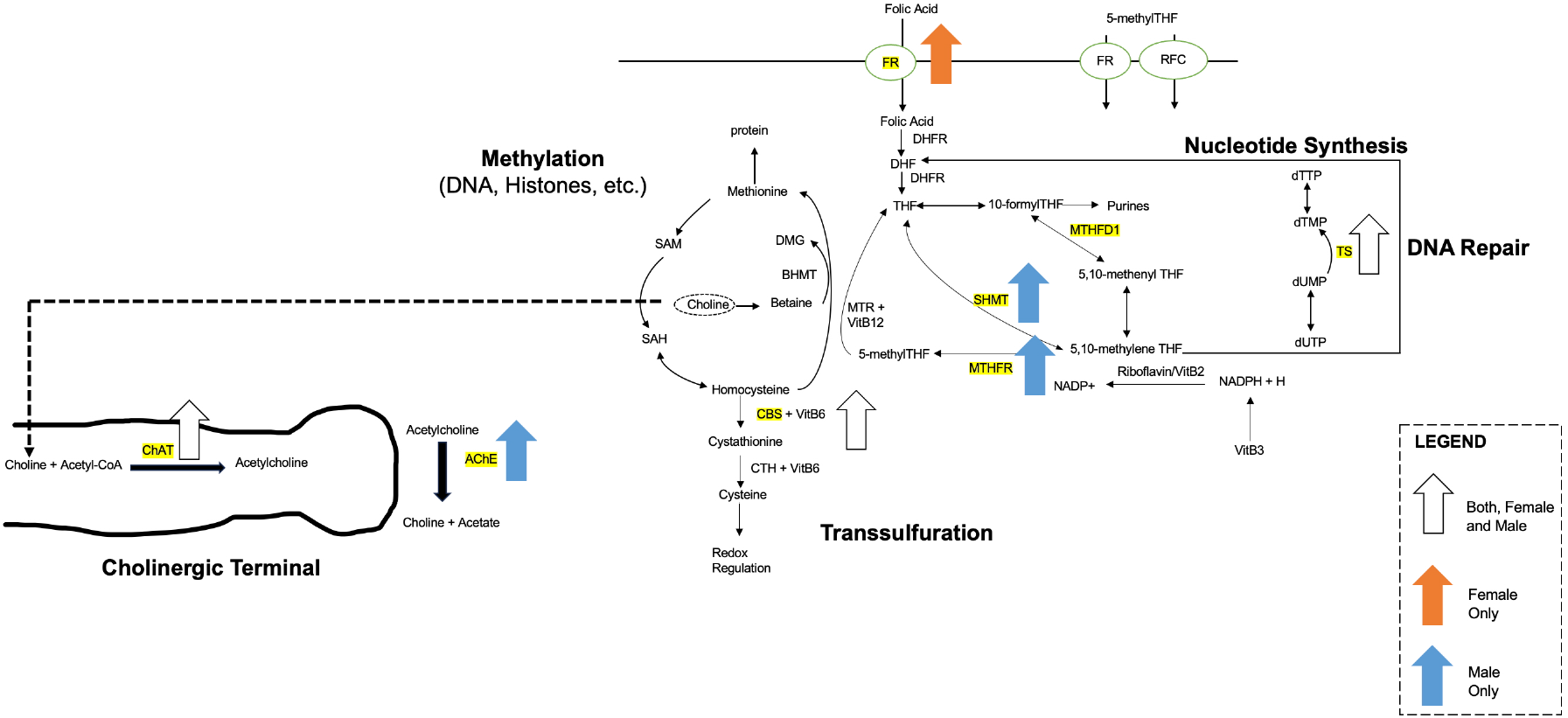

## INTRODUCTION

Ischemic stroke ranks as the second most common cause of death worldwide and predominantly affects individuals aged 65 and older^1,2^. With the changing demographics and aging of the global population, the incidence of medical emergencies like stroke is expected to increase^3^. Targeting risk factors of ischemic stroke, to reduce prevalence, is becoming more urgent. Nutrition is a modifiable risk factor for ischemic stroke^4,5^.

Elevated levels of homocysteine, a non-protein amino acid, are associated with an increased risk of cardiovascular disease, such as ischemic stroke^6,7^. B-vitamins including folic acid (vitamin B9) and vitamin B12 play crucial roles in reducing levels of homocysteine, through the one-carbon (1C) metabolism pathway^8^. In older individuals, decreased absorption of B vitamins from the gastrointestinal tract (GIT) can predispose them to ischemic stroke^9^. In addition to cofactor deficiency, the enzymes that work to metabolize homocysteine could contain inherent genetic changes. For example, methylenetetrahydrofolate reductase (MTHFR) is a critical enzyme in methylating homocysteine thereby studies have characterized the effects of a 677 C→ T genetic polymorphism in this enzyme with development of vascular disease^10^.

Stroke occurs more frequently in men than in women of the same age. However, in total, women experience more strokes than men because women have a longer life expectancy and a much higher incidence of stroke at older ages^11^. Stroke incidence in women increases significantly after menopause, suggesting that hormonal changes might play a role. The prevalence and incidence of stroke increases in postmenopausal women, leading to higher mortality rates and case fatality rates for older women, 60% of stroke fatalities are attributed to women^12^. It is believed premenopausal women are protected from stroke due to mechanisms related to sex hormones. Estrogen appears to be neuroprotective, enhancing blood flow by reducing vascular reactivity, while testosterone has opposing effects. It is also important to consider that women tend to live longer, which affects their stroke statistics^11^. There is still much to be discovered about the variations in stroke between men and women.

Though the current literature demonstrates elevated homocysteine levels with an increased risk of ischemic stroke, in animal models there is no direct comparison of 1C enzyme levels between control human brain tissue and post-mortem human ischemic stroke brain tissue. Also, currently, there is no literature evaluating the impact of sex differences on 1C enzyme levels. The aim of this study was to investigate whether ischemic stroke contributes to a change in the levels of 1C enzymes in the post-mortem brain tissue of male and female patients.

## METHODS

Human brain tissue was obtained from Banner Sun Health Research Institute Brain and Body Donation Program^16^. Study was determined to be IRB exempt as per the Midwestern University IRB Committee. Cerebral brain tissue was obtained from female and male ischemic stroke patients and healthy control subjects. Details of samples including age and comorbidities are listed in Table 1.

**Table 1.**
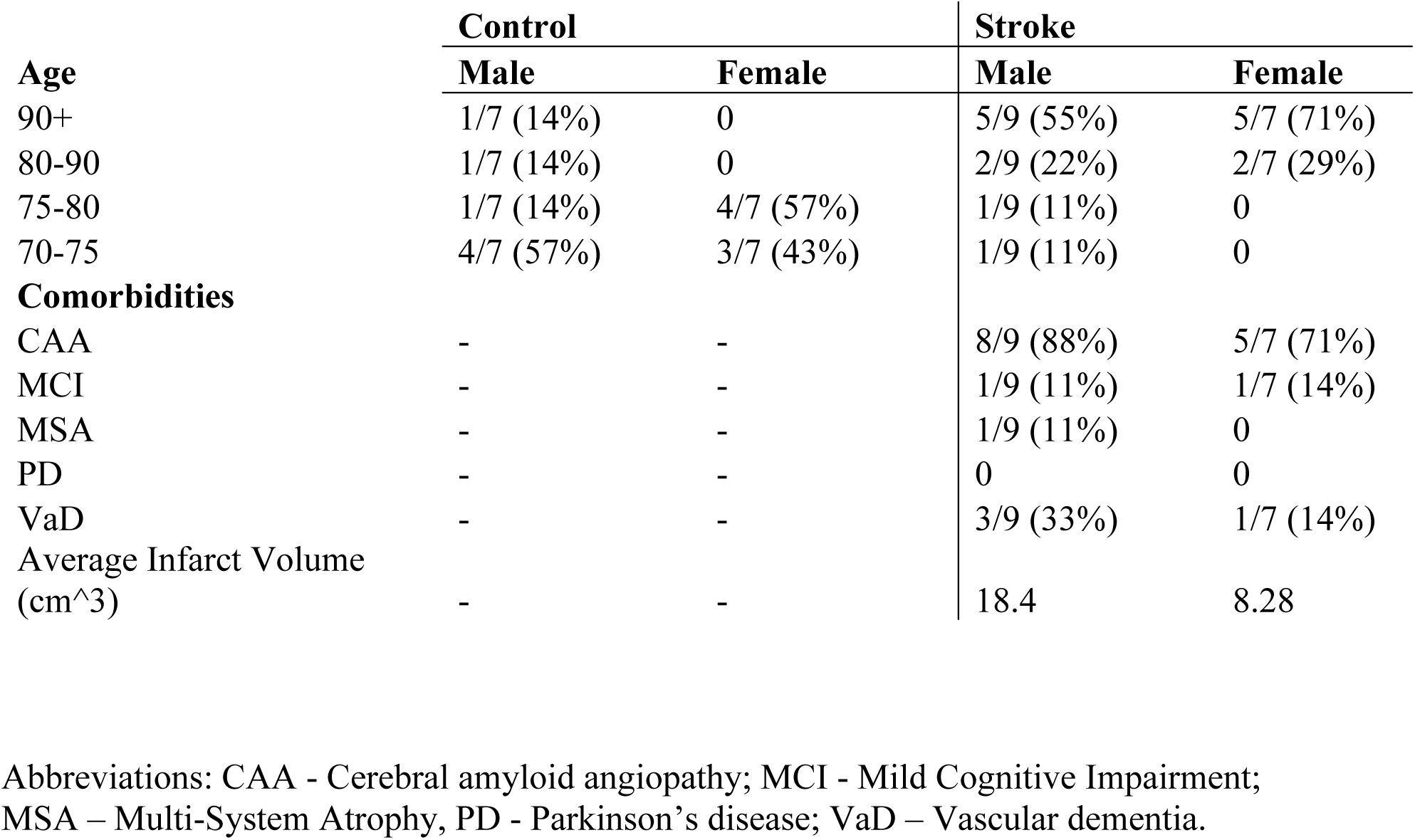
Patient sample demographics and comorbidities.

|  | **Control** |  | **Stroke** |  |
| --- | --- | --- | --- | --- |
|  | **Male** | **Female** | **Male** | **Female** |
| **Age** |  |  |  |  |
| 90+ | 1/7 (14%) | 0 | 5/9 (55%) | 5/7 (71%) |
| 80-90 | 1/7 (14%) | 0 | 2/9 (22%) | 2/7 (29%) |
| 75-80 | 1/7 (14%) | 4/7 (57%) | 1/9 (11%) | 0 |
| 70-75 | 4/7 (57%) | 3/7 (43%) | 1/9 (11%) | 0 |
| **Comorbidities** |  |  |  |  |
| CAA | - | - | 8/9 (88%) | 5/7 (71%) |
| MCI | - | - | 1/9 (11%) | 1/7 (14%) |
| MSA | - | - | 1/9 (11%) | 0 |
| PD | - | - | 0 | 0 |
| VaD | - | - | 3/9 (33%) | 1/7 (14%) |
| Average Infarct Volume<br>(cm<sup>3</sup>) | - | - | 18.4 | 8.28 |

Abbreviations: CAA - Cerebral amyloid angiopathy; MCI - Mild Cognitive Impairment; MSA – Multi-System Atrophy, PD - Parkinson’s disease; VaD – Vascular dementia.

### Immunofluorescence Experiments

To investigate 1C enzyme levels co-expression within neuronal nuclei, immunofluorescence experiments of ischemic stroke and control brain tissue was performed (**Figure 1**). The following primary antibodies were used to identify the specific protein in question for a given sample: FR (1:100, Thermo Fisher, Waltham, MA, USA, RRID: AB_2609390), MTHFR (1:100, AbCam, Boston, MA, USA, RRID: AB_2893493), SHMT (1:100, Cell Signaling, Danvers, MA, USA, catalog # 80715), TS (1:100, Cell Signaling, Danvers, MA, USA, catalog # 9045), ChAT (1:100, Millipore Sigma, Darmstadt, Germany, RRID: AB_2079751), AChE (1:100, Millipore Sigma, Darmstadt, Germany, RRID: AB_10602654) were used. All brain sections were stained with a marker for neuronal nuclei, NeuN (1:200, AbCam, Waltham, MA, USA, RRID: AB_10711040).

**Figure 1.**
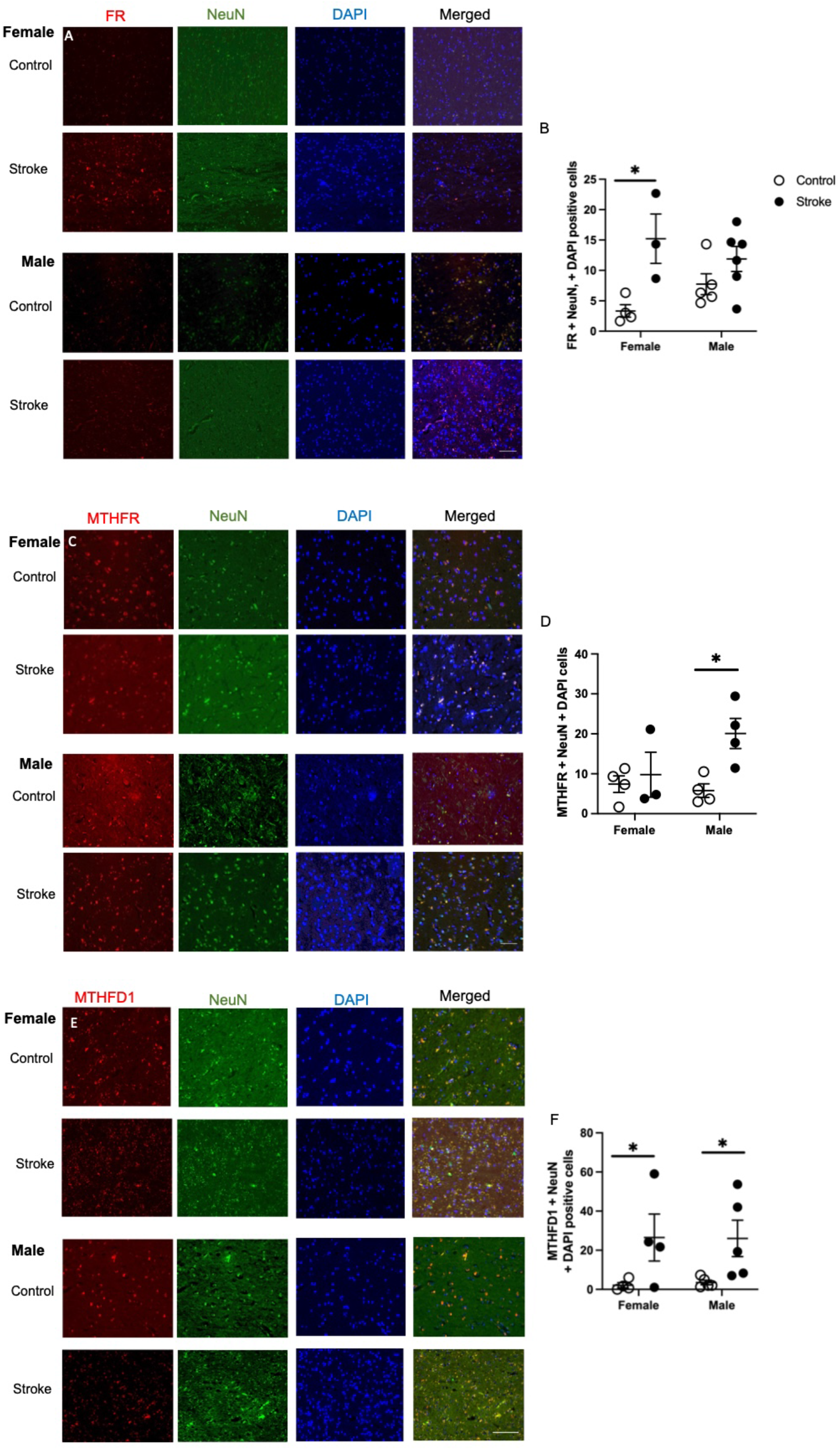
Representative images of the levels of folate receptor (FR; A), methylenetetrahydrofolate reductase (MTHFR; B), and methylenetetrahydrofolate dehydrogenase 1 (MTHFD1; E) co-stained with neuronal nuclei (NeuN) and DAPI in brain tissue of healthy aged controls and ischemic stroke patients. Quantification of FR (B), MTHFR (D), and MTHD1 (E) using a semi-quantitative method. Data represents 3 to 5 patients per group. 2-way ANOVA revealed difference between control and stroke patients for FR, MTHFR, and MTHFD1 levels. ^*^ p < 0.05, Tukey’s pairwise comparison between female control and stroke patients. Scale bar at 50 μm.

Primary antibodies were diluted in 0.5% Triton X and incubated on the brain tissue overnight at 4°C. Brain sections were incubated with secondary antibodies Alexa Fluor 488 or 555 (1:200, Cell Signaling Technologies) the following day at room temperature for 2h then stained with 4′, 6-diamidino-2phenylindole (DAPI, 1:10,000). Microscope slides were cover slipped with fluromount and stored at 4℃ until analysis.

### Immunofluorescence Analysis

The staining was visualized using the Revolve microscope (Echo, San Diego, CA, USA) and all images were collected at the magnification of 200×. We used a semi-quantitative method to assess 1C protein levels within ischemic stroke and control brain tissue. Specifically, colocalization of the primary antibody with NeuN and DAPI-labeled neurons was counted and averaged across three sections per primary antibody per subject. Using FIJI (NIH)^17^ cells were distinguished from debris by identifying a clear cell shape and intact nuclei (indicated by DAPI and NeuN) under the microscope. All cell counts were conducted by at least two individuals blinded to treatment groups.

### Statistics

GraphPad Prism 10.5.0 was used to immunofluorescence staining quantification. In GraphPad Prism a D’Agostino-Pearson normality test was performed prior to two-way ANOVA analysis when comparing the mean measurement of both sex and group (control or stroke). Significant main effects of two-way ANOVAs were followed up with Tukey’s post-hoc test to adjust for multiple comparisons. All data are presented as mean + standard error of the mean (SEM). Statistical tests were performed using a significance level of 0.05.

## RESULTS

### Increased levels of folate receptor in female stroke patients

Representative images of folate receptor neuronal staining are shown in **Figure 1A**. Ischemic stroke impacted neuronal folate receptor levels in cortical tissue (**Figure 1B**, F(_1,14_) = 12.72, p = 0.0031). Females stroke patients had more expression of the folate receptor compared to healthy controls (p = 0.021).

### Male stroke patients have increased levels of methylenetetrahydrofolate reductase (MTHFR)

Representative images of neuronal MTHFR staining are shown in **Figure 1C**. Ischemic stroke affected levels of neuronal MTHFR in cortical brain tissue (**Figure 1D**, F(_1,11_) = 12.72, p = 0.029). Male ischemic stroke patients had higher levels of MTHFR compared to healthy controls (p =0.038).

### Methylenetetrahydrofolate dehydrogenase (MTHFD1) elevated in female and male stroke patients

Representative images of neuronal MTHFD1 staining are shown in **Figure 1E**. Ischemic stroke affected levels of neuronal MTHFD1 levels in cortical brain tissue (**Figure 1F**, F (_1,14_) = 9.74, p = 0.0075). Both female (p = 0.0471) and male (p = 0.0413) had increased levels of neuronal MTHFD1 compared to healthy controls.

### Thymidylate synthase (TS) levels elevated in female and male patients

Representative images of neuronal TS staining are shown in **Figure 2A**. Ischemic stroke impacted levels of neuronal TS in cortical brain tissue (**Figure 2B**, F (_1,13_) = 22.59, p = 0.0004). Both female (p = 0.013) and male (p = 0.0453) ischemic stroke patients had higher levels of neuronal TS in brain tissue.

**Figure 2.**
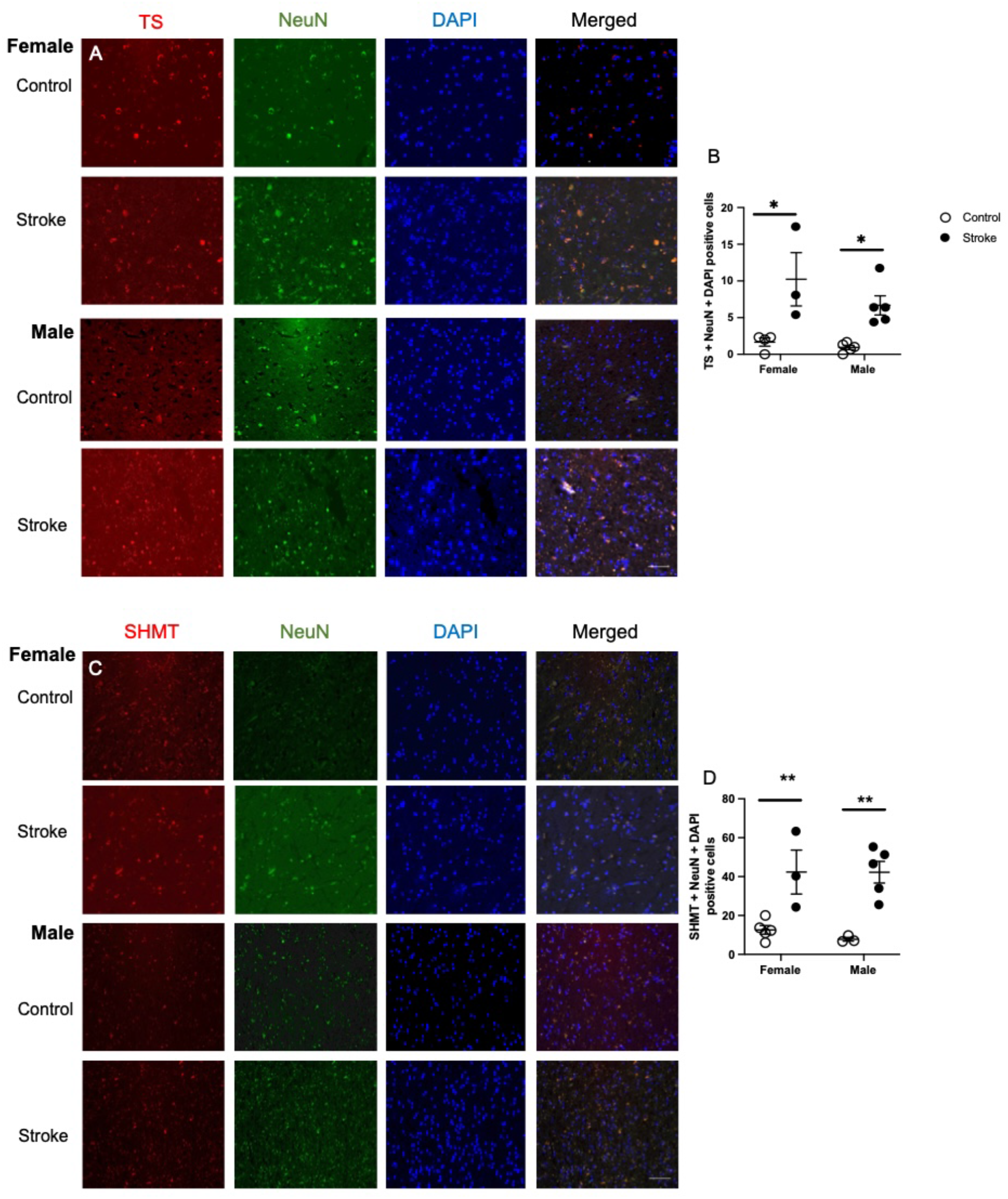
Representative images of the levels of thymidylate synthase (TS; A) and serine hydroxymethyltransferase (SHMT; B), co-stained with neuronal nuclei (NeuN) and DAPI in brain tissue of healthy aged controls and ischemic stroke patients. Quantification of TS (B) and SHMT (D) using a semi-quantitative method. Data represents 3 to 5 patients per group. 2-way ANOVA revealed difference between control and stroke patients for TS and SHMT levels. ^*^ p < 0.05, ^**^ p< 0.01, Tukey’s pairwise comparison between control and stroke male or female patient. Scale bar at 50 μm.

### Serine hydroxymethyltransferase (SHMT) levels increased in females and males

Representative images of neuronal SHMT staining are shown in **Figure 2C**. Ischemic stroke impacted levels of neuronal SHMT in cortical brain tissue (**Figure 2D**, F (_1,12_) = 9.15, p = 0.0085). Both female (p = 0.003) and male (p = 0.0012) ischemic stroke patients had higher levels of neuronal SHMT in brain tissue.

### Choline acetyltransferase (ChAT) levels impacted by stroke and gender

Representative images of neuronal ChAT staining are shown in **Figure 3A**. Ischemic stroke affected levels of neuronal ChAT levels in cortical brain tissue (**Figure 3B**, F(_1,14_) = 10.47, p = 0.0060). There was also a gender difference between female and male patients (F(_1,14_) = 5.39, p = 0.036). There was no interaction between gender and stroke (p = 0.32).

**Figure 3.**
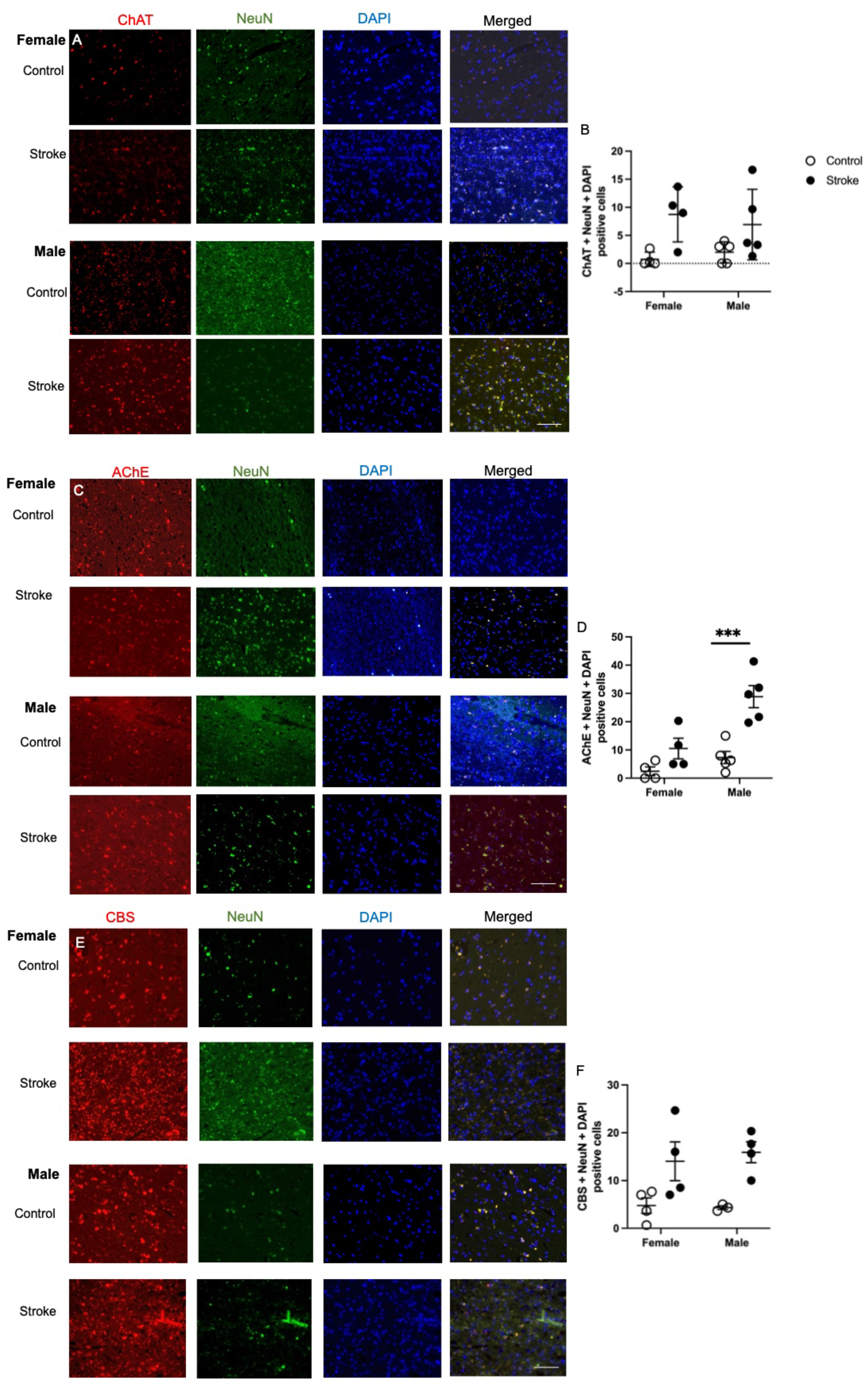
Representative images of the levels of choline acetyltransferase (ChAT; A), acetylcholinesterase (AChE; B), and Cystathionine-β-synthase (CBS; E) co-stained with neuronal nuclei (NeuN) and DAPI in brain tissue of healthy aged controls and ischemic stroke patients. Quantification of ChAT (B), AChE (D) and CBS (F) using a semi-quantitative method. Data represents 3 to 5 patients per group. 2-way ANOVA revealed difference between control and stroke patients for TS and SHMT levels. ^***^ p < 0.001, Tukey’s pairwise comparison between control and stroke male or female patient. Scale bar at 50 μm.

### Ischemic stroke and gender impact acetylcholinesterase (AChE) levels

Representative images of neuronal AChE staining are shown in **Figure 3C**. Ischemic stroke affected levels of neuronal AChE levels in cortical brain tissue (**Figure 3D**, F(_1,14_) = 23.31, p = 0.0003). There was also a gender difference between female and male patients (F(_1,14_) = 14.39, p = 0.0020) and an interaction between gender and health status (F(_1,14_) = 4.894, p = 0.0441). Male ischemic stroke patients had higher levels of neuronal AChE compared to healthy controls (p = 0.0006).

### Ischemic stroke increases levels of Cystathionine-β-synthase (CBS)

Representative images of neuronal CBS staining are shown in **Figure 3E**. Ischemic stroke affected levels of neuronal CBS levels in cortical brain tissue (**Figure 3F**, F(_1,11_) = 15.41, p = 0.0024). There was impact of gender (p = 0.90) and interaction between gender and stroke (p = 0.77).

## DISCUSSION

One-carbon (1C) metabolism plays a crucial role in various physiological processes, including biosynthesis, amino acid homeostasis, epigenetic maintenance, and redox protection. Further, current literature does indicate that elevated homocysteine is a risk factor for ischemic stroke^18,19^. However, data of 1C enzyme levels between healthy and stroke-affected human brain tissue are lacking, as well as whether there are gender differences. Our study aimed to determine whether ischemic stroke alters 1C enzyme levels by analyzing post-mortem cortical brain tissue from both male and female patients. All 1C enzymes we measured were increased in stroke patients compared to healthy controls. Interestingly, we observed sex differences. Notably, our results indicated female ischemic stroke patients had elevated expression of the folate receptor when compared to healthy controls. This suggests an increased demand of folates in the brain post-stroke in females. Male ischemic stroke patients had elevated MTHFR and AChE levels compared to healthy controls. Both female and male stroke patients showed significantly elevated levels of MTHFD1, SHMT, and TS compared to healthy controls. Our data demonstrates the impact of sex on 1C enzyme levels following ischemic stroke in cortical brain tissue. These observed alterations in 1C enzyme expression may correspond with the established sex-specific differences in stroke presentation, risk factors, and metabolic responses post-stroke.

Between sexes, presenting symptoms of stroke differ, with women reporting atypical stroke symptoms at a higher prevalence as compared to men^20–22^.Women report headaches, consciousness or mental status changes, and coma, while men typically report paresis/hemiparesis and focal visual disturbances. Women usually present with more severe stroke symptoms and are 4 to 5 years older than men at onset^20^. Further, young men have a much higher risk of stroke compared to premenopausal women. However, this trend flips after menopause, leading to the incidence of stroke in older women being twice that of older men. A reversal often attributed to the loss of hormone production from the ovaries that coincides with menopause^21^. Congruently, an increased risk of stroke has been indicated in women with later age at menarche, >15 years of age, and a short reproductive life span, ≤ 36 years of age. Overall, a short reproductive life span for women is associated with an increased risk of nonfatal cardiovascular disease events, such as stroke, in midlife^23^.

Sex differences in metabolism after ischemic stroke are increasingly apparent but remain unexplored. Limited studies suggest that females exhibit greater changes in amino acid and lipid metabolism after stroke, including higher levels of neurotoxic kynurenine metabolites, which are linked to worse post-stroke outcomes. Further, one preclinical study showed that Huang-Lian-Jie-Du decoction improved neurological function after stroke in both male and female rat models, with greater benefit in females, potentially due to sex-specific differences in metabolites linked to oxidative stress regulation^24^. These findings begin to highlight how sex differences in 1C metabolism mirror the differing outcomes seen post-stroke between sexes. The present study has added to this body of literature demonstrating sex differences in 1C enzyme levels in elderly male and female stroke patients.

One-carbon metabolism with folate support is essential for nucleotide synthesis, homocysteine remethylation, and numerous biosynthetic processes critical for growth and development. Thus, disruptions in folate metabolism can impair cellular proliferation, influence epigenetic regulation, and increase the risk of congenital disabilities or disease^19^. Studies show men exhibit higher plasma concentrations of choline, betaine, and homocysteine and an increased likelihood of folate deficiency. In the present study we report increased levels of neuronal levels of AChE in male stroke patients and increased levels of ChAT in both stroke females and males. AChE is an enzyme involved in breaking down acetylcholine at the synapse and ChAT is an enzyme that synthesizes it. Previous work from our group has shown that choline compensates in the brain during elevated levels of homocysteine, as a result of deficiencies in folate metabolism, either genetic^25^ or dietary^26^. It is well documented that aging results in increased levels of homocysteine^27^, indicating the need for increased levels of AChE and ChAT in the entirety of the aging population.

The importance of 1C metabolism can also be seen by the increases in MTHFD1, SHMT, TS, and CBS in both stroke females and males in the present study. MTHFD1 encodes a key enzyme in folate-mediated 1C metabolism, essential for nucleotide synthesis and methylation, with disruptions linked to metabolic imbalance and cognitive deficits^28^. Its increase post-stroke, likely reflects a compensatory response post-stroke, aimed at boosting one-carbon metabolism to support DNA repair, methylation, and reduce harmful homocysteine through enhanced folate-driven reactions.

SHMT is a vitamin B_6_-dependent enzyme that catalyzes the reversible conversion of serine and tetrahydrofolate into glycine and 5,10-methylene-THF, supplying one-carbon units essential for nucleotide synthesis, methylation, and redox balance^29^. Its increase in the present study aligns with the literature that found ischemic stroke patients exhibit significantly higher SHMT1 promoter methylation compared to healthy controls, which is strongly associated with elevated plasma homocysteine levels. This suggests that stroke triggers epigenetic changes that suppress SHMT1 expression, prompting compensatory metabolic responses to normalize folate-dependent one-carbon metabolism and manage toxic homocysteine accumulation^30^.

TS catalyzes the conversion of deoxyuridine monophosphate (dUMP) to deoxythymidine monophosphate (dTMP) using 5,10-methylene-THF as a methyl donor, which provides the only *de novo* source of thymine nucleotides essential for DNA synthesis and repair^31^. The findings in the present study conform with accepted idea that after ischemic stroke, specific TS gene polymorphisms are more common and are associated with increased TS expression, which likely enhances DNA repair and one-carbon metabolism to counteract tissue damage and elevated homocysteine levels^32^. Thus, stroke patients may exhibit upregulated TS activity as a compensatory response to vascular injury and metabolic stress.

CBS catalyzes the first and rate-limiting step of the transsulfuration pathway, combining homocysteine and serine to form cystathionine. This is a crucial process for detoxifying neurotoxic homocysteine and producing cysteine for glutathione synthesis to support antioxidant defenses. After ischemic stroke, elevated homocysteine becomes neurotoxic and pro-inflammatory. As described in the literature and seen in the present study, CBS is upregulated post-stroke to convert homocysteine into cystathionine, aiding in detoxification and supporting antioxidant defense via glutathione synthesis, thus serving as a crucial compensatory mechanism against oxidative stress in the damaged brain^19^. These changes in 1C metabolism highlight the effects and underscore the complications of ischemic stroke seen in stroke patients.

In women estrogen upregulates phosphatidylethanolamine N-methyltransferase, enhancing phosphatidylcholine synthesis and influencing 1C flux. These differences in metabolism and hormones contribute to sex-specific methylation patterns^33^. In men, choline and homocysteine levels are associated with specific epigenetic modification that regulates DNA repair and oncogenic risk^33,34^. Meanwhile, choline and vitamin B12 correlate with protective epigenetic modification essential for genome stability and hematopoiesis in women^25^. Further, mathematical modeling indicates estrogen-driven upregulation of phosphatidylethanolamine methyltransferase in women significantly elevates choline and betaine concentrations. This increase enriches tissue-level activation of cystathionine β-synthase, which is the primary enzyme responsible for homocysteine clearance. As a result, females exhibit lower plasma homocysteine levels compared to males^35^. Overall, this previous work and our study highlights how sex-specific regulation of 1C metabolism shapes not only nutrient status, but also disease risk, developmental outcomes, and epigenetic programming.

One-carbon (1C) metabolism plays an important role in the generation of S-adenosylmethionine, a global methyl donor^36^. Changes in DNA methylation after ischemic stroke have been reported in preclinical models and patients ranging from hyper to hypomethylation ^37,38^. This data suggests that DNA methylation has a role for inducing stroke pathologies and/or managing recovery mechanisms after disease onset. Changes in methylation of pathophysiological events after stroke including excitotoxicity, oxidative stress, mitochondrial dysfunction, blood-brain barrier (BBB) disruption, apoptosis and inflammation have been reported ^37,39^. Furthermore, there have been recovery changes including neurogenesis, angiogenesis, gliogenesis, axon growth, and synaptic plasticity ^37^. Understanding the implication of methylation pattern changes post-stroke is an area that needs more attention.

In the present study, stroke females had significantly increased levels of folate receptor possibly coinciding with the neuroprotective nature of estrogen. Estrogen does this by triggering the rapid retention of kinases like PI3K/Akt and ERK, leading to enhanced cell survival signaling and reduced oxidative stress in peri-infarct tissues. These same signaling pathways are also involved in upregulating nutrient transport systems like folate metabolism during repair from damage, i.e., ischemic stroke^40^. Further, these sex differences in 1C metabolism may also explain the significantly increased levels of MTHFR found in males. The literature suggests that testosterone boosts MTHFR activity, while estradiol suppresses it, indicating that males inherently have higher MTHFR activity due to their androgen levels^35^. This suggests that after a stroke, male patients are more likely to upregulate MTHFR in response to ischemic stress and homocysteine accumulation. In contrast, females, influenced by estrogen, maintain more stable MTHFR levels.

This is the first study to investigate protein levels of 1C enzymes after ischemic stroke in cortical tissue of aged patients. Our study was limited to the cerebral cortex as well as measuring 1C enzyme levels in neuronal cells. Expanding to other areas of the brain and investigating the impact of stroke on other cell types (e.g., glial cells) are needed to grain a comprehensive understanding of the impact ischemic stroke has on 1C enzyme levels. Furthermore, expanding data collection on homocysteine levels as well as other 1C metabolites would add to the data set. A comprehensive understanding of how ischemic stroke impacts and interactions with 1C will enable future therapeutic developments with a precision medicine focus. Furthermore, the changes in DNA methylation post-stroke and link to 1C are an exciting opportunity for investigation.

## Acknowledgements

We are grateful to the Banner Sun Health Research Institute Brain and Body Donation Program of Sun City, Arizona for the provision of human biological materials. The Brain and Body Donation Program has been supported by the National Institute of Neurological Disorders and Stroke (U24 NS072026 National Brain and Tissue Resource for Parkinson’s Disease and Related Disorders), the National Institute on Aging (P30 AG19610 and P30AG072980, Arizona Alzheimer’s Disease Center), the Arizona Department of Health Services (contract 211002, Arizona Alzheimer’s Research Center), the Arizona Biomedical Research Commission (contracts 4001, 0011, 05-901 and 1001 to the Arizona Parkinson’s Disease Consortium) and the Michael J. Fox Foundation for Parkinson’s Research.

GEM was awarded the Mentored Professional Enrichment Experience award from Southern Illinois University School of Medicine.

This work was supported by the American Heart Association, Grant Number 20AIREA35050015 to NMJ.

